# CRISPRCasTyper: An automated tool for the identification, annotation and classification of CRISPR-Cas loci

**DOI:** 10.1101/2020.05.15.097824

**Authors:** Jakob Russel, Rafael Pinilla-Redondo, David Mayo-Muñoz, Shiraz A. Shah, Søren J. Sørensen

## Abstract

CRISPR-Cas loci encode for highly diversified prokaryotic adaptive defense systems that have recently become popular for their applications in gene editing and beyond. The increasing demand for bioinformatic tools that systematically detect and classify CRISPR-Cas systems has been largely challenged by their complex dynamic nature and rapidly expanding classification. Here, we developed CRISPRCasTyper, a new automated software tool with improved capabilities for identifying and typing CRISPR arrays and *cas* loci across prokaryotic sequences, based on the latest classification and nomenclature (39 subtypes/variants) (Makarova et al. 2020; Pinilla-Redondo et al. 2019). As a novel feature, CRISPRCasTyper uses a machine learning approach to subtype CRISPR arrays based on the sequences of the direct repeats. This allows the typing of orphan and distant arrays which, for example, are commonly observed in fragmented metagenomic assemblies. Furthermore, the tool provides a graphical output, where CRISPRs and *cas* operon arrangements are visualized in the form of colored gene maps, thus aiding annotation of partial and novel systems through synteny. Moreover, CRISPRCasTyper can resolve hybrid CRISPR-Cas systems and detect loci spanning the ends of sequences with a circular topology, such as complete genomes and plasmids. CRISPRCasTyper was benchmarked against a manually curated set of 31 subtypes/variants with a median accuracy of 98.6%. Altogether, we present an up-to-date and freely available software pipeline for significantly improved automated predictions of CRISPR-Cas loci across genomic sequences.

**Implementation:** CRISPRCasTyper is available through conda and PyPi under the MIT license (https://github.com/Russel88/CRISPRCasTyper), and is also available as a web server (http://cctyper.crispr.dk).

## Introduction

CRISPR-Cas systems constitute a group of bacterial and archaeal adaptive immune systems that have garnered much attention in recent years due to their promising biotechnological applications (Pickar-Oliver and Gersbach 2019; Barrangou and Doudna 2016). These systems are composed of two main components: 1) the CRISPR array, a chromosomal memory bank of sequences derived from previous infecting genetic parasites, and 2) operon(s) of CRISPR-associated (*cas*) genes encoding the proteins required during the three stages of immunity (adaptation, processing, and interference). For further details on the mechanisms driving the CRISPR-Cas immune response, we refer readers to recent reviews (Hille et al. 2018; Mohanraju et al. 2016; Jackson et al. 2017).

Like all other prokaryotic defense systems, CRISPR-Cas loci evolve rapidly in a constant arms race with their mobile genetic element foes (Hampton, Watson, and Fineran 2020). The resultant evolutionary tension has led to a remarkable diversification of CRISPR-Cas systems, which, together with the apparently frequent exchange of components and lack of a universal marker gene across systems (Koonin and Makarova 2019), greatly challenges the development of a unified classification scheme. Accordingly, classification efforts have relied on a multi-faceted approach that jointly takes into consideration the architectural organization of CRISPR-Cas loci, the presence/absence of certain Cas components, and sequence similarities of genes (Makarova et al. 2015). Broadly, the current classification contemplates two major classes, Class 1 and Class 2, that either rely on heteromeric multi-protein effector complexes or single multi-domain effector proteins, respectively (Makarova et al. 2020). In the next hierarchical level, there are six types (I, III and IV for Class 1; and II, V and VI for Class 2), each of which contain several subtypes and multiple variants. While recent years have seen an extraordinary expansion in the classification of newly discovered systems, the current classification is predicted to be nearly complete at the “type” level (Makarova, Wolf, and Koonin 2018; Makarova et al. 2020). For a summary of the state of the art classification and nomenclature, we refer the readers to recent comprehensive reviews and articles (Makarova et al. 2020; Pinilla-Redondo et al. 2019).

The systematic efforts to classify novel CRISPR-Cas systems have run parallel to those aiming their automated prediction across genomic sequences. Although systematic CRISPR-Cas identification pipelines have been developed (Crawley, Henriksen, and Barrangou 2018; Couvin et al. 2018; Lange et al. 2013), their sensitivity below the type level is generally inadequate. Furthermore, the discovery of novel systems has been occurring at a breakneck pace, rendering older classification software obsolete. Additionally, many CRISPR-Cas loci are complex, comprising hybrid *cas* cassettes or share arrays between different Cas types. For instance, recent work has shown that around 40% of CRISPR-Cas loci show atypical organizations, where orphan CRISPR arrays and *cas* operons are common, as well as hybrid loci resulting from the associations of different co-occurring systems within genomes (e.g. distinct types of interference modules and one shared adaptation cassette) (Bernheim et al. 2020). However, the CRISPR-Cas prediction tools published so far largely lack the formalism required to handle such complexity.

Here we present CRISPRCasTyper, a new tool that can accurately identify and annotate CRISPR and *cas* loci automatically based on the newest classification (Makarova et al. 2020; Pinilla-Redondo et al. 2019). Besides classifying *cas* operons, CRISPRCasTyper can also accurately assign subtypes to CRISPR arrays based on the sequence composition of the consensus repeat. We also provide the first benchmark of automated classification of CRISPR-Cas loci on a manually curated dataset across all known subtypes, which exemplifies the strengths of CRISPRCasTyper.

## Software description

CRISPRCasTyper identifies *cas* operons and associated CRISPR arrays from an input fasta-formatted DNA sequence (Fig. 1A). CRISPRCasTyper searches for *cas* and other genes functionally linked to CRISPR-Cas systems with HMMER3 (Eddy 2009) against 680 Hidden Markov Models (HMMs). Matches to Class 2 effectors (*cas9, cas12, cas13*) and the III-E gRAMP fusion protein are filtered by specific E-value and coverage cutoffs optimized specifically for each effector. The remaining HMM matches are filtered by overall cutoffs (see Methods and Materials for details). Adjacent *cas* and accessory genes are then joined into operons; inclusion of a gene in the operon is based solely on synteny. These operons are then typed based on a scoring scheme (see Methods and Materials for details).

**Figure 1.**
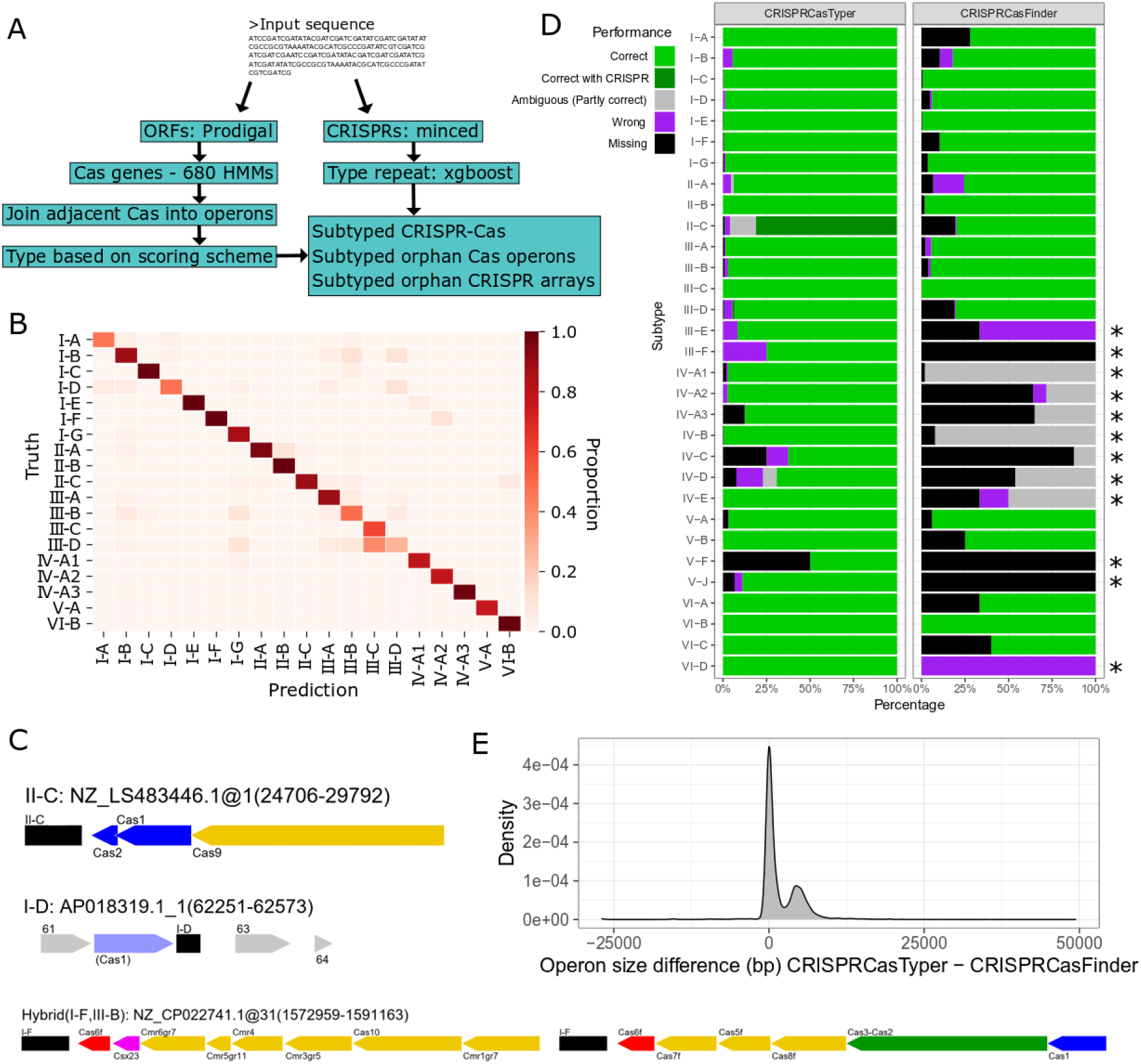
**A)** CRISPRCasTyper workflow. **B)** Prediction matrix of CRISPR repeat typer against an unseen test dataset. Only subtypes with at least 20 repeats were included in the model. Counts are normalized per row totals. **C)** Examples of graphical outputs from CRISPRCasTyper. The predicted subtype associated with a repeat sequence is written above the array, colored in black (top). Interference module in yellow (except Cas6), adaptation module in blue, Cas6 in red, and accessory genes in purple. Unknown genes (grey) and genes with low-quality matches (same color scheme in lighter shade) can be added to the plots, also around orphan CRISPR arrays (middle). Furthermore, CRISPRCasTyper resolves hybrid systems (bottom). **D)** Performance of CRISPRCasTyper versus CRISPRCasFinder on manually curated CRISPR-Cas systems. ‘Correct with CRISPR’ denotes loci which were resolved using subtype prediction based on the CRISPR repeat sequence. Asterisks (*) denote subtypes not included in CRISPRCasFinder; CRISPRCasFinder includes type IV without subtype prediction, and these are denoted as ‘Ambiguous’. **E)** Density plot of differences in operon sizes between CRISPRCasTyper and CRISPRCasFinder.

CRISPRCasTyper includes the following 39 subtypes/variants: I-A, I-B, I-C, I-D, I-E, I-F, I-G, II-A, II-B, II-C, III-A, III-B, III-C, III-D, III-E, III-F, V-A, V-B, V-C, V-D, V-E, V-F, V-G, V-H, V-I, V-J, VI-A, VI-B, VI-C, VI-D (Makarova et al. 2020), IV-A1, IV-A2, IV-A3, IV-B, IV-C, IV-D, IV-E (Pinilla-Redondo et al. 2019), and transposon-associated V-K (Strecker et al. 2019) and I-F (Klompe et al. 2019).

To aid in resolving ambiguous CRISPR-Cas operons, and to subtype distant and orphan CRISPR arrays, we created a CRISPR repeat classification model. We used gradient boosting decision trees fitted to counts of canonical tetramers (Chen and Guestrin 2016, see Methods and Materials for details). Only subtypes with at least 20 known repeats were included. The classifier has a median accuracy across the 19 included subtypes of 89% on an unseen test dataset (Fig. 1B). Furthermore, the web server includes an additional model, which is automatically re-trained monthly using subtyped repeats crowdsourced from the inputs from web server users. This novel feature ensures that the accuracy of the tool increases over time and with usage of the platform, as well as its ability to recognize previously undetectable subtypes/variants. As of writing, this model includes more than ∼34k repeats from non-redundant genomes, in addition to the ∼6k repeats in the manually curated set. This model also includes the subtypes IV-D, IV-E, V-B, V-F, V-J, VI-A, and VI-D (Fig. S1), and has a median per-subtype accuracy of 84%, mainly drawn down by the rare subtypes III-C, IV-D, and V-B.

As an additional feature, CRISPRCasTyper automatically draws gene maps to enable visualization of the operonic structure (Fig. 1C). These gene maps can be expanded to include HMM matches below the inclusion thresholds, which could aid the discovery of diversified *cas* components or accessory genes, partial and novel CRISPR-Cas variants/subtypes, especially around orphan CRISPR arrays.

Moreover, CRISPRCasTyper can resolve loci spanning the ends of circular sequences, such that loci are not erroneously split in two due to the linear representation of circular sequences. Furthermore, the percentage completion of both the interference and adaptation modules of each detected operon are provided in the output. Finally, CRISPRCasTyper runs in less than a minute on a typical genome (2-6 Mbp) using 4 threads, and in less than 10 minutes on a deep metagenome assembly (60-100 Mbp) using 20 threads.

## Benchmark

CRISPRCasTyper was compared with the widely used CRISPRCasFinder (Couvin et al. 2018) using the manually curated CRISPR-Cas loci from Makarova et al. 2020 (Makarova et al. 2020), the newest type IV classification from Pinilla-Redondo et al. 2019 (Pinilla-Redondo et al. 2019) and manually curated III-E loci (see Methods and Materials). We find that CRISPRCasTyper outperforms CRISPRCasFinder at identifying 17 subtypes and is equally accurate in the prediction of the remaining 2 subtypes included in CRISPRCasFinder (Fig. 1D) (Couvin et al. 2018). Across these 19 subtypes, CRISPRCasTyper had a median accuracy of 99.5%, while that of CRISPRCasFinder was 93.9%. The median accuracy of CRISPRCasTyper on all 31 subtypes (12 lacking in CRISPRCasFinder) was 98.6%. Furthermore, CRISPRCasTyper often provides a more complete *cas* operon identification (Fig. 1E). An example is a type III-B operon in an *Acidilobus saccharovorans* genome (NC_014374.1), in which CRISPRCasFinder identifies *cmr4, cmr5*, and *cmr6*, whereas CRISPRCasTyper finds 9 additional genes, including *cas1, cas2, cas4, cas10*, and *cas6*.

Both CRISPRCasTyper and CRISPRCasFinder found few false positives (28 (0.4%) and 24 (0.4%), respectively), with a large bias towards VI-B (10 and 13, respectively). Interestingly, several of these seem to be true positives, which have been missed in the curated dataset; 9 of the VI-B operons have adjacent CRISPR arrays whose repeat sequence is predicted by CRISPRCasTyper to be VI-B associated (Table S1). V-F is especially challenging due to the similarity of its effector to transposases; we chose a conservative approach, which identifies as many as possible without finding false positives. Many V-F are missed with this approach, but some can be found using the graphical output if they are adjacent to CRISPR arrays. When the V-F subtype is more clearly defined, improved HMMs might solve this problem.

## Methods

### CRISPRCasTyper

Open reading frames are called with prodigal v2.6.3 (Hyatt et al. 2010). Protein-profile alignments are performed with HMMER3 (Eddy 2009). HMM matches are filtered in a two-step process. All single-effector genes (*cas9, cas12, cas13, gRAMP*) are filtered with specific cutoffs (see Table S2). The remaining Cas proteins are filtered with an E-value cutoff of 0.01, and sequence and HMM coverage both of 30%. The single-effector cutoffs were set by running a grid search across the curated set with coverages between 5% and 95% with a step size of 5%, and E-values from 10e-5 to 1e-150 with a step size of the exponent of 5. The subtypes with no representative in the curated set were given the following cutoffs: E-value: 1e-5, sequence coverage: 90%, HMM coverage: 25%. *Cas* genes are grouped into operons based on synteny. By default, no more than 3 unannotated genes can separate known *cas* genes to be considered part of a single operon.

The operons are then typed based on a scoring scheme where each Cas HMM within it has a score between 0 and 4 for each subtype. The scoring scheme was built such that mandatory HMMs give a score of 2 and accessory genes a score of 1. HMMs specific to a subtype adds 2 to the score. Forbidden HMMs give a score of -4. The subtype that obtains the highest score is then assigned to the operon if the score is at least 6 and there are at least 3 different *cas* genes. Operons with at least 6 *cas* genes and two or more types with a score of at least 6 and at least one specific HMM are denoted as ‘Hybrid’ systems. Operons falling outside these cutoffs are annotated as ‘Putative’ unless one of the genes is a Class 2 effector or the III-E gRAMP fusion protein. Non-hybrid operons with multiple equally scoring subtypes are annotated as ‘Ambiguous’.

To resolve problems of ambiguity, a score of 0.1 was added to mandatory and specific I-F HMMs, such that typing defaults to type I-F, unless TniQ is found which will change the type to I-F_T, transposon-associated I-F. Similarly for IV-A2, which is distinguished from IV-A1 and IV-A3 by the absence of Csf1. Typing therefore defaults to IV-A2 unless a Csf1 is found or Csf2 or Csf4 is specific to IV-A1 or IV-A3. Similarly with III-A and III-F, for which the scores are designed to default to III-A unless SSgr11 is detected. As Cas12j and Cas12k are so similar the typing defaults to V-J unless transposon-associated TniQ, TnsB, and TnsC are found.

### RepeatTyper

A curated set of subtyped repeat sequences was created by predicting CRISPR arrays with minced v0.4.2 (https://github.com/ctSkennerton/minced, Bland et al. 2007) in the curated datasets (Makarova et. al. 2020; Pinilla-Redondo et. al. 2019). Consensus repeats from all arrays within 1kbp to a *cas* operon were included. This resulted in a total of 5838 subtyped repeat sequences. Only subtypes with at least 20 repeat sequences were included in the model. For each repeat sequence all canonical tetramers were counted, and these 136 features were the input for our model. The sequences were split in 70% training data, used to train the model and choose parameters, and 30% testing data, used as an unseen dataset to evaluate the accuracy of the final model. We used xgboost v1.0.2 with multi:softprob objective evaluated with mlogloss and 3-fold cross-validation across a grid of max-depth={4, 6, 8}, subsample={0.6, 0.8, 1}, and colsample_bytree={0.6, 0.8, 1}. The models were run with a learning rate of 0.3 for 100 boosting rounds, but with 10 early stopping rounds. The remaining parameters were defaults. The script for training this model is part of CRISPRCasTyper, such that users can easily re-run the model on their own repeat-set and/or with other parameters. The accuracy for each subtype was calculated as percent correct predictions on the test set. The adjusted accuracy was then the average accuracy across all subtypes, such that subtypes with many repeats were not inflating the accuracy.

On the web server, consensus repeats from novel loci are automatically included in the model. Novelty is based on position of loci in the sequence, the subtype prediction, and the repeat sequence. This updated set was supplemented with subtyped repeats from Ensembl bacterial and archaeal genomes (Yates et al. 2020), GTDB (Parks et al. 2018), and NCBI metagenomes (ftp://ftp.ncbi.nlm.nih.gov/genomes/genbank/metagenomes/). As of writing this article, the model includes 40717 subtyped repeats across 37 subtypes/variants; 26 of these with at least 20 repeat sequences. This updated model is automatically re-trained each month with the same parameters as above except with max-depth={6, 8, 10}.

### Hidden Markov Models

CRISPRCasTyper includes 680 HMMs. The largest share was built using hmmbuild from HMMER3 v3.2.1 (Eddy 2009) on the multiple alignments provided by Makarova et al. 2020 (Makarova et al. 2020), excluding the consensus sequence. Some HMMs were obtained from CRISPRCasFinder 2.0.2. The V-K specific HMMs and the I-F associated TniQ HMM were built from multiple alignments created with MUSCLE v3.8.1551 (Edgar 2004), based on previous studies (Strecker et al. 2019; Klompe et al. 2019). Type IV HMMs were obtained from Pinillia-Redondo et. al. 2019.

### Benchmark

All non-type IV CRISPR-Cas loci from Makarova et al. 2020, available at ftp://ftp.ncbi.nih.gov/pub/wolf/_suppl/CRISPRclass19/, were included. Multi-systems and partial systems were excluded from the benchmark. Type IV loci were obtained from Pinilla-Redondo et. al. 2019 to include the newest classification. We further included a curated set of III-E loci (Table S3). For CRISPRCasTyper, all predicted non-putative systems were included. For CRISPRCasFinder, all systems, including partial systems, were included. As Makarova et al. 2020 does not include V-K, the few V-J operons which were predicted by CRISPRCasTyper to be V-K were labelled as correct classifications. For determining false positives only operons not overlapping with any loci, including partial and multi-systems, were counted. CRISPRCasFinder version 4.2.17 with CasFinder 2.0.2 was used for the benchmark. CRISPRCasTyper version 1.0.0 was used for the benchmark.

### Data availability

The scoring table, all HMM profiles, the filtering cut-offs, the repeat typer model, and the completion files are available in the data directory at the CRISPRCasTyper github: https://github.com/Russel88/CRISPRCasTyper/tree/master/data. The updated repeat typer models are available at http://mibi.galaxy.bio.ku.dk/russel/repeattyper/.

### Web server

The web server was built on the jobson framework (https://github.com/adamkewley/jobson), with a modified UI available at https://github.com/Russel88/jobson. The web server also includes the possibility to submit a RefSeq/Genbank accession, which will download the corresponding nucleotide sequence through the entrez-direct command line tool.

## Acknowledgments

We would like to thank Sarah Camara for testing the software and providing valuable feedback. J.R and S.A.S were supported by the Novo Nordisk Foundation. R.P-R was financed by the Independent Research Fund Denmark, InTrans Project [#8022-00322B]. D.M-M was supported by a University of Otago Doctoral Scholarship. Part of the data analysis was performed on Computerome, the Danish National Computer for Life Sciences.

## Supplementary Information

**Figure S1.**
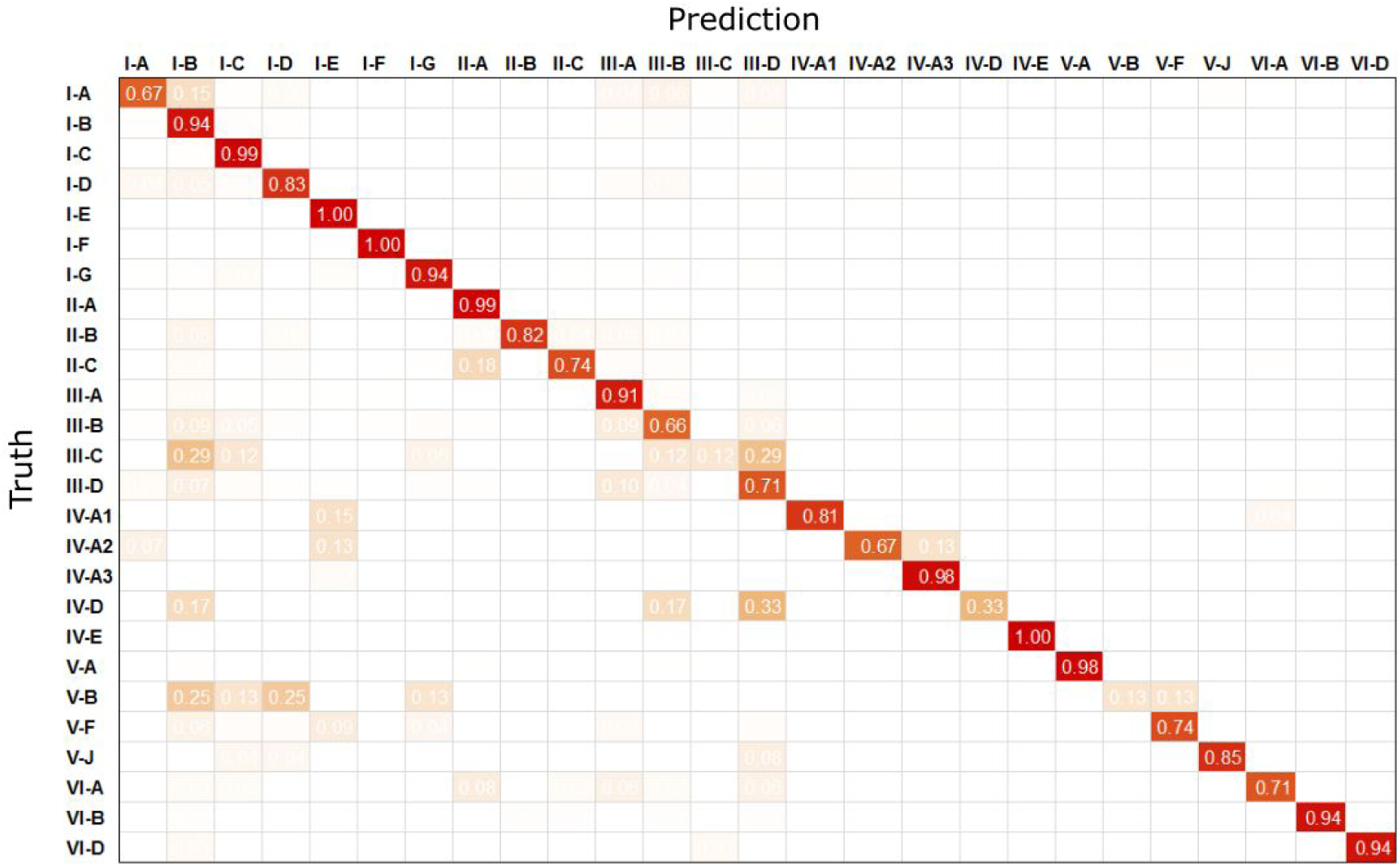
Prediction matrix of CRISPR repeat typer against an unseen test dataset using the updated repeat set of 40k repeat sequences. Counts are normalized per row totals, such that the diagonals contain the accuracy for each subtype.

**Table S1.** False positive operons from CRISPRCasTyper. Several have CRISPRs adjacent, and the predicted subtype of these CRISPRs match the CRISPRCasTyper predictions. * The subtype for the repeat sequence from the V-J on NZ_AP014815.1 could not be predicted with the curated repeat set, but was predicted to be V-J with the updated repeat set. § This I-E loci is likely part of the I-E loci in the end of this sequence.

| Nucleotide | Subtype | Start | End | CRISPR | CRISPR Subtype |
| --- | --- | --- | --- | --- | --- |
| NC_014377.1 | I-A | 1179284 | 1183097 | No |  |
| NC_017768.1 | I-A | 838904 | 841348 | No |  |
| NZ_CP022385.1 | I-A | 1996653 | 2002119 | No |  |
| NZ_CP031218.1 | I-B | 1882001 | 1884693 | No |  |
| NZ_CP019794.1 | I-C | 4407864 | 4411147 | Yes | I-C |
| NC_008750.1 | I-D | 2011041 | 2015216 | No |  |
| NC_009438.1 | I-D | 2601641 | 2605816 | No |  |
| NC_014248.1 | I-D | 4303670 | 4309589 | No |  |
| NC_017566.1 | I-D | 2602859 | 2607034 | No |  |
| NC_015499.1 | I-E | 1031980 | 1036308 | No |  |
| NZ_CP032329.1 | I-E § | 3 | 7027 | Yes | I-E |
| NZ_CP032099.1 | II-B | 801907 | 802326 | No |  |
| NC_015519.1 | III-A | 3080 | 4920 | No |  |
| NC_019954.2 | III-A | 3199 | 5039 | No |  |
| NZ_CP021838.1 | III-B | 2057 | 4497 | No |  |
| NC_002950.2 | VI-B | 1244197 | 1248317 | Yes | VI-B |
| NC_010729.1 | VI-B | 1410847 | 1414967 | Yes | VI-B |
| NC_014734.1 | VI-B | 3133284 | 3136748 | No |  |
| NC_016001.1 | VI-B | 2545026 | 2549183 | Yes | VI-B |
| NZ_CP011995.1 | VI-B | 920122 | 924242 | Yes | VI-B |
| NZ_CP012889.1 | VI-B | 1409775 | 1413895 | Yes | VI-B |
| NZ_CP024591.1 | VI-B | 209480 | 213600 | Yes | VI-B |
| NZ_CP024595.1 | VI-B | 1078280 | 1082400 | Yes | VI-B |
| NZ_CP025930.1 | VI-B | 1411695 | 1415815 | Yes | VI-B |
| NZ_CP025932.1 | VI-B | 1167344 | 1171464 | Yes | VI-B |
| NC_019678.1 | V-J | 6308425 | 6308943 | No |  |
| NZ_AP014815.1 | V-J | 2551679 | 2553529 | Yes | Unknown (V-J)* |
| NZ_CP026681.1 | V-J | 5056609 | 5057136 | No |  |

**Table S2.**
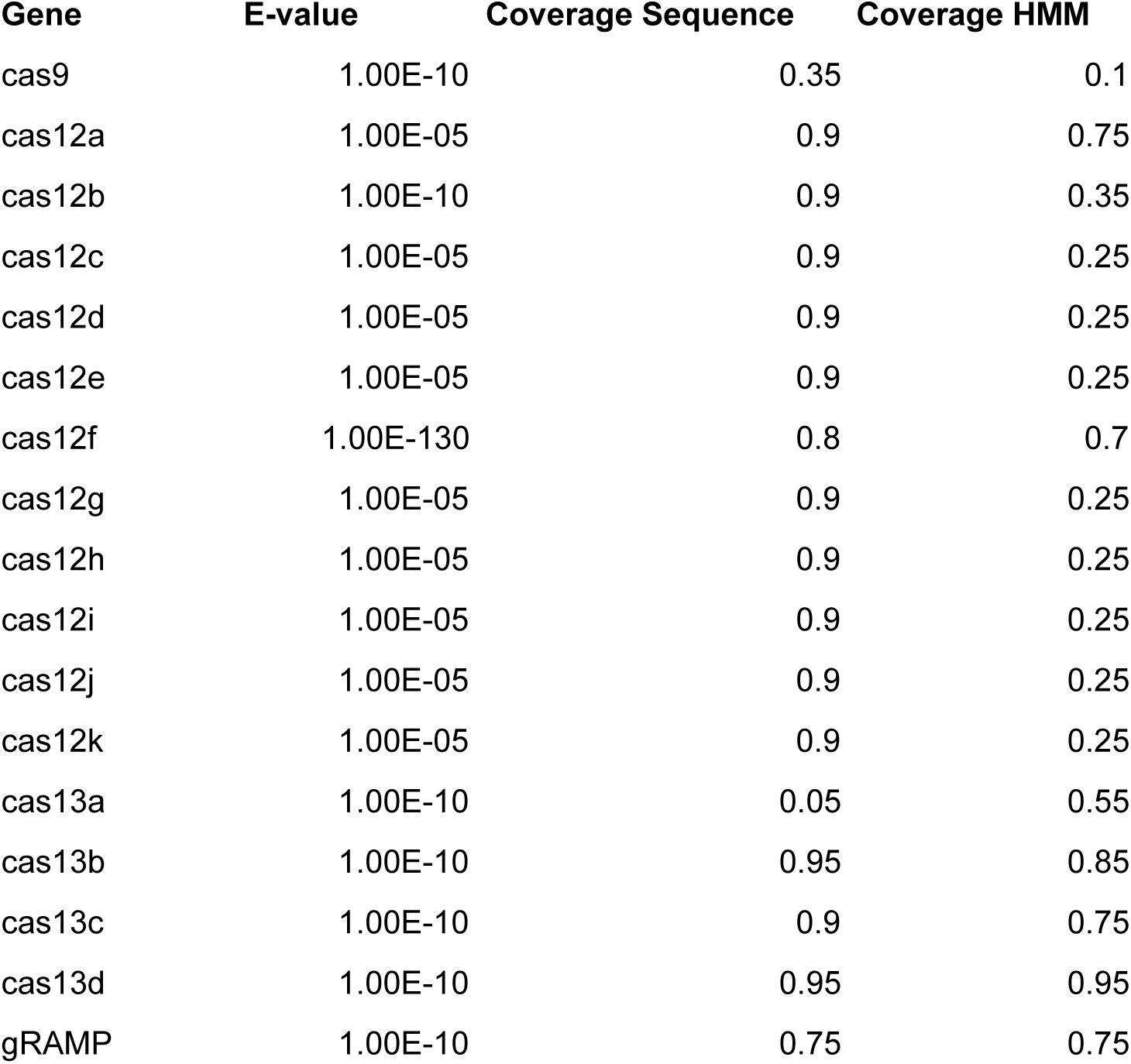
Specific cutoffs used for filtering single-effector *cas* genes.

| Gene | E-value | Coverage Sequence | Coverage HMM |
| --- | --- | --- | --- |
| cas9 | 1.00E-10 | 0.35 | 0.1 |
| cas12a | 1.00E-05 | 0.9 | 0.75 |
| cas12b | 1.00E-10 | 0.9 | 0.35 |
| cas12c | 1.00E-05 | 0.9 | 0.25 |
| cas12d | 1.00E-05 | 0.9 | 0.25 |
| cas12e | 1.00E-05 | 0.9 | 0.25 |
| cas12f | 1.00E-130 | 0.8 | 0.7 |
| cas12g | 1.00E-05 | 0.9 | 0.25 |
| cas12h | 1.00E-05 | 0.9 | 0.25 |
| cas12i | 1.00E-05 | 0.9 | 0.25 |
| cas12j | 1.00E-05 | 0.9 | 0.25 |
| cas12k | 1.00E-05 | 0.9 | 0.25 |
| cas13a | 1.00E-10 | 0.05 | 0.55 |
| cas13b | 1.00E-10 | 0.95 | 0.85 |
| cas13c | 1.00E-10 | 0.9 | 0.75 |
| cas13d | 1.00E-10 | 0.95 | 0.95 |
| gRAMP | 1.00E-10 | 0.75 | 0.75 |

**Table S3.** Accession numbers of III-E gRAMP proteins. No adjacent CRISPR could be determined (NA) in short contigs or with a *gRAMP* gene at the end of a sequence.

| Nucleotide | Gene | CRISPR |
| --- | --- | --- |
| JRYO01000185.1 | KHE91659.1 | Yes |
| NZ_BAFH01000003.1 | WP_007220849.1 | Yes |
| MVRP01000104.1 | OPY65763.1 | Yes |
| MGTA01000040.1 | OGR07205.1 | Yes |
| NZ_BEXT01000001.1 | WP_124327589.1 | Yes |
| LAQJ01000233.1 | KKO18793.1 | NA |
| NBMK01000156.1 | OQY58162.1 | NA |
| QMMU01000439.1 | RLC14096.1 | NA |
| QMMU01000323.1 | RLC15988.1 | NA |
| QMMU01000137.1 | RLC19861.1 | NA |
| JPDT01001326.1 | KPA14974.1 | No |
| NZ_BEXT01000001.1 | WP_124327589.1 | No |

## Notes

### Competing Interest Statement

The authors have declared no competing interest.

